# When is a fish stock collapsed?

**DOI:** 10.1101/329979

**Authors:** J Yletyinen, WE Butler, G Ottersen, KH Andersen, S Bonanomi, FK Diekert, C Folke, M Lindegren, MC Nordström, A Richter, L Rogers, G Romagnoni, B Weigel, JD Whittington, T Blenckner, NC Stenseth

**Author notes:** These authors contributed equally to this work (J Yletyinen, WE Butler). Corresponding authors, these authors contributed equally to this work.

## Abstract

Marine fish stock collapses are a major concern for scientists and society due to the potentially severe impacts on ecosystem resilience, food security and livelihoods. Yet the general state of harvested fish populations has proven difficult to summarize, and the actual occurrence rate of stock collapses remains unclear. We have carried out a literature review and multi-stock analysis to show that numerous definitions exist for classifying stocks as collapsed, and that the classification of a stock’s status is sensitive to changes in the collapse definition’s formulation. We suggest that the lack of a unified definition has contributed to contrasting perceptions on the state of fish stocks. Therefore, we comprehensively define what constitutes a fish stock collapse and provide a time-series based method for collapse detection. Unlike existing definitions, our definition is process-based, because it links together three important phases of collapse events: the abrupt decline, an ensuing period of prolonged depletion, and potential recovery. Furthermore, these phases are specified in terms of population turnover. Through systematic evaluation, our definition can accurately distinguish collapses from less severe depletions or natural fluctuations for stocks with diverse life histories, helping identify the stocks in greatest need of reparatory measures. Our study advocates the consistent use of definitions to limit both alarmist and conservative narratives on the state of fish stocks, and to promote cooperation between conservation and fisheries scientists. This will facilitate clear and accurate communication of science to both the public and to policy-makers to ensure healthy fish stocks in the future.

## BACKGROUND

The effects of overfishing have expanded from a predominantly local to global scale in the past century. The present state of fisheries has proven difficult to summarize without generalizations affected by personal perceptions and data preferences (1,2). Consequently, scientific literature contains numerous diagnoses of global trends. During the past 15-20 years, high profile articles have reported that fisheries have caused a general decline in fish stocks worldwide (3–10), but numerous replies contested these claims (11–14). In particular, the projection of a universal fish stock collapse by 2048 (15) heated the scientific debate, especially between conservation and fisheries scientists (discussed in 2, 16–18), and triggered considerable media attention to the reportedly disastrous state of the world’s fish stocks.

In 2006, Hilborn raised the issue that high-impact journals publish papers on the decline and collapse of fisheries for their publicity value rather than scientific merit (19). This is a noteworthy concern, as words used to report scientific results may increasingly be chosen based on marketability rather than the content of the findings (i.e., may be driven by publication pressure) (20). Furthermore, it has been suggested that environmental challenges in general are neglected by the media and politics if they are not easily adapted to sensation-driven news (21). On the other hand, while the glass half-empty view on fish stock health may produce alarmist narratives, the glass half-full view can allow continued fishing pressure even when stocks are at persistently low levels. Such an overly optimistic view likely increased the severity of major stock crashes, such as the Canadian Northern cod (*Gadus morhua*) around 1990 (22) and the Norwegian spring spawning herring (*Clupea harengus*) in the late 1960s (23). Moreover, prolonged heavy fishing of already over-exploited stocks is likely to delay rebuilding and substantially increase the uncertainty in recovery time (24).

A precise and robust way to classify fish stock status is not only essential for scientific rigor and interdisciplinary cooperation, but also because ‘state of the resource’ classifications may trigger political responses with potentially far-reaching biological, social, and economic consequences. In this study, we focus on collapsed fish stocks as the most extreme classification of resource status. Collapses of targeted fish stocks are of specific interest to scientists and the general public due to the high associated risks for livelihoods, ecosystem resilience, food security and cultural meaning (4,15,25,26). At the present time of unprecedented anthropogenic impacts on the environment, the risks of ecological collapses are increasingly discussed (27,28,37–41,29–36), albeit not always thoroughly conceptualized or defined (33). Collapses have been reported in marine fisheries in many parts of the world, both in relation to fish stocks and to social or social-ecological elements, such as profitability or a specific livelihood (10,42–48).

To move towards a shared understanding of what constitutes a collapsed fish stock we suggest establishing a common definition of “collapse”. We first show that a large number of different definitions exist, and suggest that the lack of a single standard definition has led to confusion and conflicting views regarding the state of fish stocks. Ambiguity over how collapse thresholds are generated illustrates the lack of a common understanding of what constitutes a fish stock collapse. Together with different interpretations of catch and biomass data – especially between fisheries and conservation scientists (2,12) – this has led to contrasting collapse classifications and added noise and disagreement to scientific debate. To address this matter, we propose a new, quantitative definition to identify fish stock collapses using time series data. Importantly, we do not wish to just add one more method to the already extensive list, nor do we intend to provide means for detecting a collapse in advance, as we consider this to be another field of research (statistical early warning indicators of population collapse, e.g., 36, 49, 50). Instead, we want to provide a comprehensive quantitative collapse definition that can be applied to a large number of important fish stocks in a consistent manner to facilitate between-stock comparisons.

## METHODS

We reviewed scientific literature from 1984 to 2015 in ecology, fisheries science, and economics. Comparable to snowball sampling, we started with the most well-established collapse articles and expanded the article selection to collapse papers cited by or citing these articles. The review was conducted with the purpose of finding the range of how a fish stock or fishery collapse is defined, not to collect all existing collapse definitions. Hence, when we had acquired approximately 20 definitions from 82 references, we considered the sample adequate and finished the review. We included social science articles in the review, but no explicit fish stock collapse definitions were found in that field.

We extracted time-series data for 20 marine fish stocks from stock assessment databases (51,52), reports (53) and peer-reviewed articles (54,55). Taken as a whole, the selection of fish stocks represents a diverse array of population dynamic trends and life-histories (Supplementary Materials Table S2). Many of the selected stocks have been classified as collapsed, whilst others display trends that would intuitively suggest that a collapse has occurred, making for a suitable database to explore the consequences of various collapse definitions. For some stocks (e.g. eastern Baltic cod) the most recent time-series contained insufficient data, therefore older stock assessments were used to facilitate between-definition comparisons. The set of stocks comprises 11 Atlantic cod stocks, chosen as this is an extensively studied species that in many ways is a keystone indicator of the health of North Atlantic ecosystems. We supplemented these stocks with nine further stocks to capture between-species life-history variation (generation times range from 1-25 years) and additional types of population trend.

Generation time estimates for each stock were taken from the literature (Supplementary Materials Table S2). If suitable values were not found, the stock’s age-at-maturity or mean age were used as generation time (and thus population turnover) surrogates. For the West Greenland cod stock, a suitable value was not found, therefore we used an arbitrary value of 5 years in accordance with previous studies that have evaluated IUCN’s extinction criteria on gadoids (56).

Each definition was coded in the R programming language and applied to each of the stocks. Estimates of the spawning stock biomass and total biomass that can sustain maximum sustainable yield (SSB_msy_ and TB_msy_ respectively) were not readily available for many stocks. For SSB_msy_, we assumed it was twice B_pa_, a rule that was found to provide reasonable estimates by Froese *et al.* (57). For TB_msy_ and MSY, estimates were taken from a Bayesian state-space Schaefer model (58), providing that the time series of TB and catches were of comparable length. The minimum number of stocks that could be applied to any definition was 15 (mean = 18), whilst the minimum number of definitions that could be applied to any stock were 12/16, 3/6 and 7/8 for SSB, TB and catch data respectively.

The output comprised all the relevant information characterizing each definition (set parameters and type of population measure), the information describing the collapse (reference biomass and threshold values, their pertaining years, and the magnitude and rate of declines), and a number of collapse metrics. These included the number of years in a collapsed, a potentially recovered or a non-collapsed state, the duration of the collapsed state, and the stock’s present status (i.e. at the end of the time-series). This gives a broad overview of how each definition captures both the immediacy of a collapse event (“abrupt decline”) and the impaired production of a collapsed state (“prolonged depletion”).

## RESULTS

Twenty different time-series-based stock level definitions were found in the literature review (Supplementary Materials Table S1). Most definitions capture a state of depletion by setting a threshold below which the stock is classified as collapsed. We identify four weaknesses in the existing definitions: 1) the definitions often lack temporal context, which can lead to classifying stocks that naturally fluctuate or gradually decline as collapsed (Figure 1A, 1B); 2) the justification for setting thresholds is often unclear or missing, and there is little numerical consistency between definitions (e.g., the percent decline from maximum historical biomass required to set a threshold varies from 1% to 25% [Supplementary Materials Table S1]); 3) definitions are related to study-specific reference values (e.g. maximum historical biomass), yet the arbitrary historical condition used as a reference point is often subject to data availability and what individual scientists perceive to be healthy fish stocks (59); 4) few definitions were formulated in a way that can account for life-history variation between stocks, which is an essential consideration for comparing status across fish stocks.

**Figure 1.**
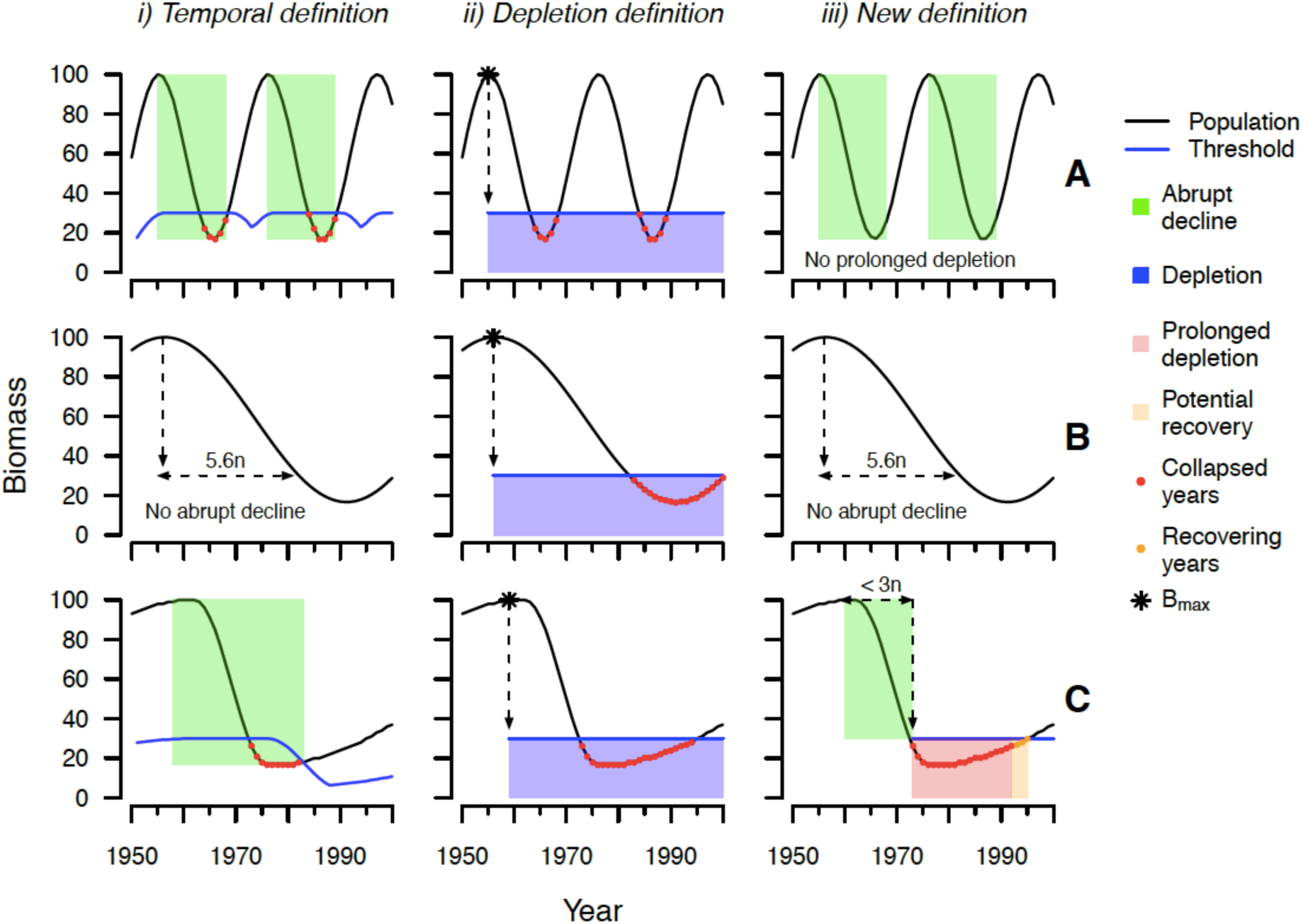
Three different collapse criteria (columns i, ii and iii) are applied to three hypothetical stocks (rows A, B and C) to demonstrate how the formulation of a collapse criteria can result in misleading classifications of a stock’s state. Stock A fluctuates in a cyclic manner, stock B gradually depletes over a long period of time, stock C rapidly declines and exhibits prolonged depletion following the decline. The first definition (i) classifies a year as collapsed if a 70 % decline occurs within three generations (i.e., it captures an abrupt decline). This can misclassify stocks with cyclic dynamics (i, A) because there is no consideration of prolonged depletion. The second definition (ii) classifies a year as collapsed if the stock’s biomass falls below 30 % of the historical maximum (B_max_). This definition misclassifies a stock that gradually depletes (B, ii) because the rate of decline is not considered, and it misclassifies a stock with cyclic dynamics (A, ii) because it does not consider prolonged depletion. The proposed definition (iii) classifies a stock as collapsed if a decline of 70 % within 3 generations is immediately followed by a period of prolonged depletion where biomass remains below the threshold for a generation. By considering the abrupt decline and prolonged depletion as an interlinked process, the proposed definition can filter out natural fluctuations (A, iii) and gradual depletions (B, iii) from more drastic collapses (C, iii).

The number of collapse definitions found in our literature review indicates that there is still no consensus regarding the choice of analytical approach used to reliably classify the status of a fish stock as collapsed. Our analysis shows that collapse definitions are highly sensitive to small changes in parameter values, such as the threshold level, and to the overall formulation of collapse definition. As an example, seven of the thirteen biomass-based definitions categorized the Irish cod stock as collapsed (Supplementary Materials Appendix A). Conversely, out of 20 stocks included in the analysis, only the Norwegian spring spawning herring, Nothern cod (2J3KL) and North Sea autumn spawning herring stocks were classified as collapsed by all applicable definitions.

The definition we propose views a collapse as an abrupt change to an undesired state. It consists of two sequential, interlinked criteria that capture the collapse process: an abrupt decline, and an ensuing period of prolonged depletion. Indicative of impaired production, prolonged depletion is an undesired state for a fish stock. Each criteria is defined as follows:

1. Abrupt decline: a stock’s adult biomass declines by 70% within a maximum of three generations or 10 years.
2. Prolonged depletion: the stock’s mean adult biomass over a succeeding generation remains below the 70% abrupt decline.

Therefore, when applied to a time-series of adult biomass (spawning stock biomass, SSB), a stock has collapsed if an abrupt decline (1) is immediately (i.e., the year at the bottom of the decline is evaluated) followed by a prolonged depletion (2). For any year *i,* the mean biomass over a succeeding generation 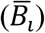 is simply a left-aligned average with an averaging window of one generation (*n*),

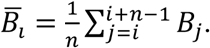

If both criteria are met, a threshold T_B_ is set at 0.3B_t_, where B_t_ is the stock’s biomass at the beginning of the abrupt decline (Fig. 1C). Subsequently, all years following the abrupt decline are evaluated for prolonged depletion (criteria 2). If a year’s mean adult biomass over a succeeding generation is below or equal to the threshold 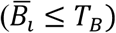, the stock remains in a collapsed state, whereas if it is above the threshold 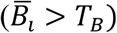, the stock is exhibiting potential recovery from the collapsed state (Fig. 1C).

If an abrupt decline is not immediately followed by a prolonged depletion, we classify the event as a temporary fluctuation instead of a collapse (Fig. 1A) because the decline is temporary (i.e., less than a generation time, thus production is not impaired). If the timespan for a 70% decline is greater than a maximum of three generations or 10 years, the event is classifed as a “gradual depletion” because the decline is not abrupt (Fig 1B). Inclusion of the abrupt decline criteria links the definition to the common use of “collapse” as “*a sudden, complete fall”* in both ecology and semantics. This distinguishes it from a less severe “decline” (“*becoming smaller, fewer or less”*, 60, 61) by specification of a threshold and a temporal window within which the decline must occur (but see 62), thus ensuring the decline is both large and abrupt. Importantly, the use of temporal window negates the shifting baseline syndrome (59) because the reference biomass is not specified in advance (e.g., maximum historical biomass, which could be significantly affected by the length of the time series). For example, the West of Scotland cod stock fell below the 10% historical maximum total biomass threshold, but not below the equivalent 90% *abrupt* decline threshold (Supplementary Materials Appendix A). This is because a 90% decline is not recorded within three generations (or 10 years) of the historical maximum in 1981, so the maximum historical total biomass is an inappropriate baseline for considering this particular stock as collapsed. The exclusion of gradually depleting stocks may seem too cautionary, but it draws from the quantitative values of IUCN (International Union for Conservation of Nature) criteria that are developed through wide consultation (63).

Previous studies (56,64) have highlighted that IUCN’s decline-rate thresholds can contribute to the detection of fish stocks most in need of restoration measures due to negative associations between decline- and recovery-rates, and through the inclusion of intrinsic life-history traits (65). We focus on the 70% threshold, but tested all three IUCN threshold values, namely 70%, 80% and 90%, in our analyses. Although we focus on the 70% threshold, the final threshold parameter should result from a scientific assessment recommending a particular global target, including policymakers’ views on what is realistically achievable, and with consideration of future ecosystem variation (66). Once a threshold is set, the baseline condition is defined in relation to the year at which the abrupt fall started. For instance, the Eastern Baltic Sea cod stock has collapsed in relation to its biomass in 1981 (Supplementary Materials Appendix A). The proposed definition is thus based on conservation thresholds, while consideration for recovery potential allows recognition of successful fisheries management. This aspect aims to unify the views of conservation science and fisheries science (16,67).

To critically evaluate our definition, we compare it with the 20 existing definitions (Supplementary Materials Table S1) to a set of 20 marine fish stocks with diverse life-histories and population dynamics (Figure 2 and Supplementary Materials Appendix A). The comparison illustrates that collapse events can be distinguished from natural fluctuations remarkably well by considering fish stock collapse as a multiphase process quantified with species/stock-specific generation times (Figures 1 and 2). For instance, Barents Sea capelin (*Mallotus villosus*) (Supplementary Materials Appendix A) is a naturally fluctuating stock classified as frequently collapsed according to many definitions, whereas our definition identifies collapse for only a small number of these years. For the 80% and 90% thresholds, our definition categorises 13% and 4% of years as collapsed respectively, as opposed to the equivalent IUCN definitions which categorise 37% and 22% of years as collapsed. Figures 3 and 4 compares the performance of our definition to the existing collapse definitions across the 20 stocks. The proportion of collapses detected (with a 70% threshold) is higher with our process-based definition than with most existing definitions. Furthermore, our definition highlights that most of the collapsed stocks (12 out of 17) have potentially recovered, with some stocks even exceeding the baseline reference biomass. This agrees well with previous recovery assessments of many of the included stocks (68).

**Figure 2.**
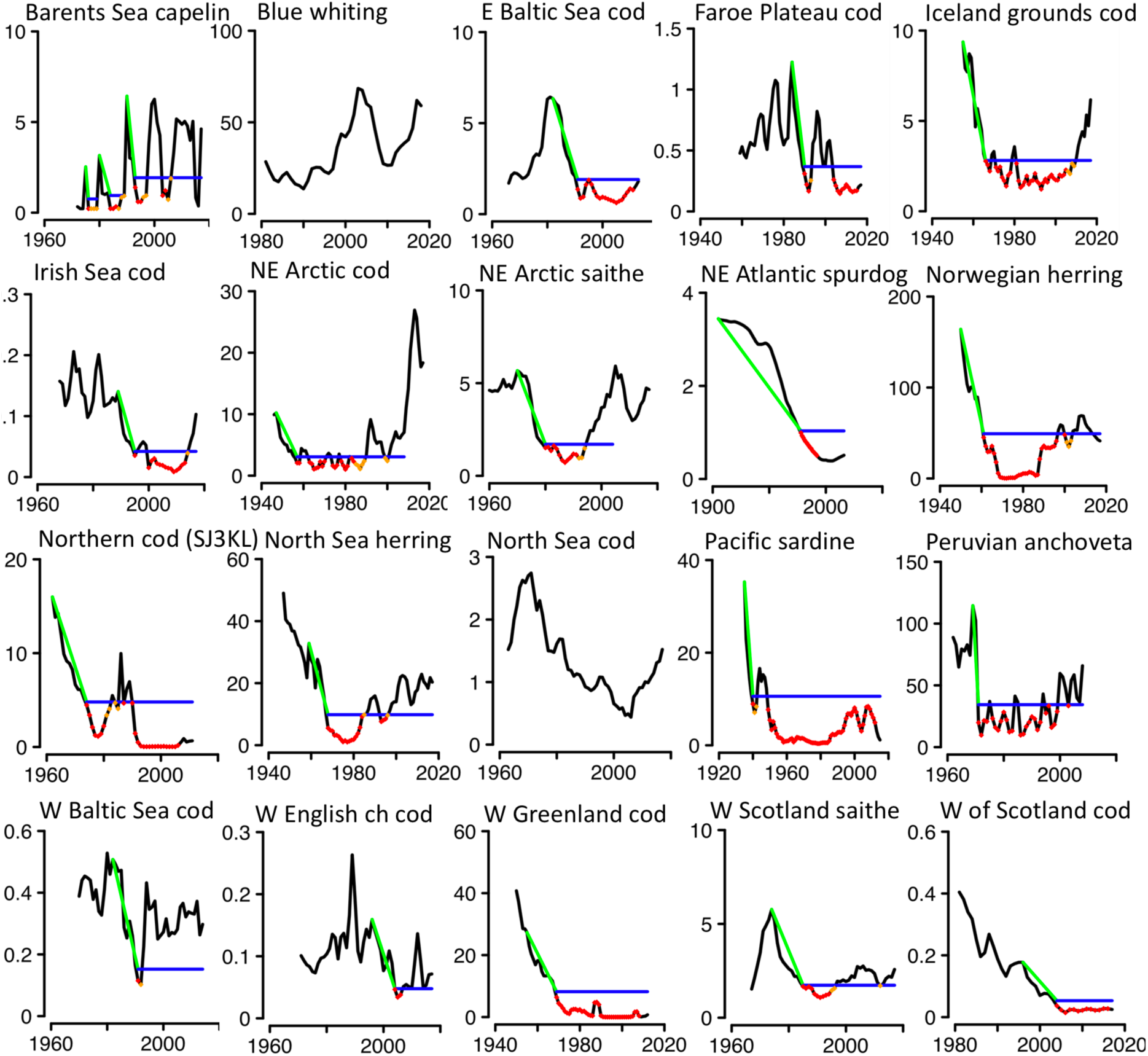
Fish stocks included in the study. The selected stocks capture a diverse array of population dynamic trends and life-histories, chosen for testing the collapse definition on a variety of fish stock dynamics. The units on the y-axis of the time series graphs are 100,000 metric tonnes; see Figure 1 for colour legend. See Supplementary Materials Appendix A for more detailed stock graphs. The following abbreviations are used. W: west, E: east, NE: northeast, and for the stocks: North Sea herring: North Sea autumn spawning herring, W Scotland saithe: North Sea, Rockall and West Scotland, Skagerrak and Kattegat saithe, North Sea cod: North Sea, Eastern English Channel, Skagerrak cod, Norwegian herring: Norwegian spring-spawning herring, W English ch cod: Western English Channel and Southern Celtic Sea cod. West Greenland cod graph is in this figure based on total biomass data due to lack of SSB data.

**Figure 3.**
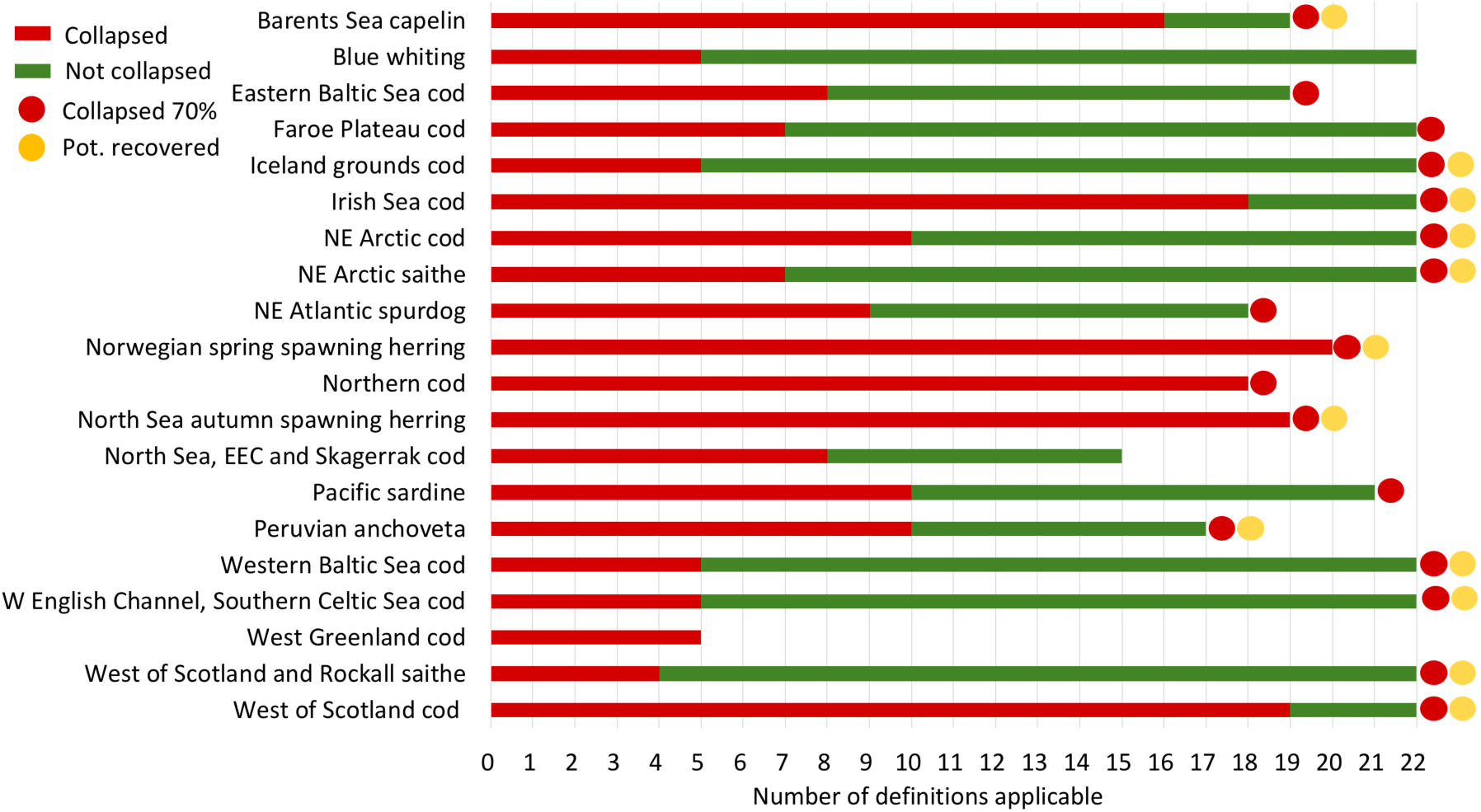
Summary of the analysis results showing the number of collapse definitions that classify stocks as collapsed vs non-collapsed. The lengths of the bars next to each stock indicate the number of existing definitions that were applicable (ranging from 6 to 18 definitions due to the various data needs of the different definitions). The dots at the ends of the bars show the status of each stock using our new proposed definition with 70 % threshold: no dot indicates that the stock has not been classified as collapsed; a red dot indicates the stock has collapsed; a subsequent yellow dot indicates a potentially recovered stock. NE: northeast, EEC: Eastern English Channel, W: western.

**Figure 4.**
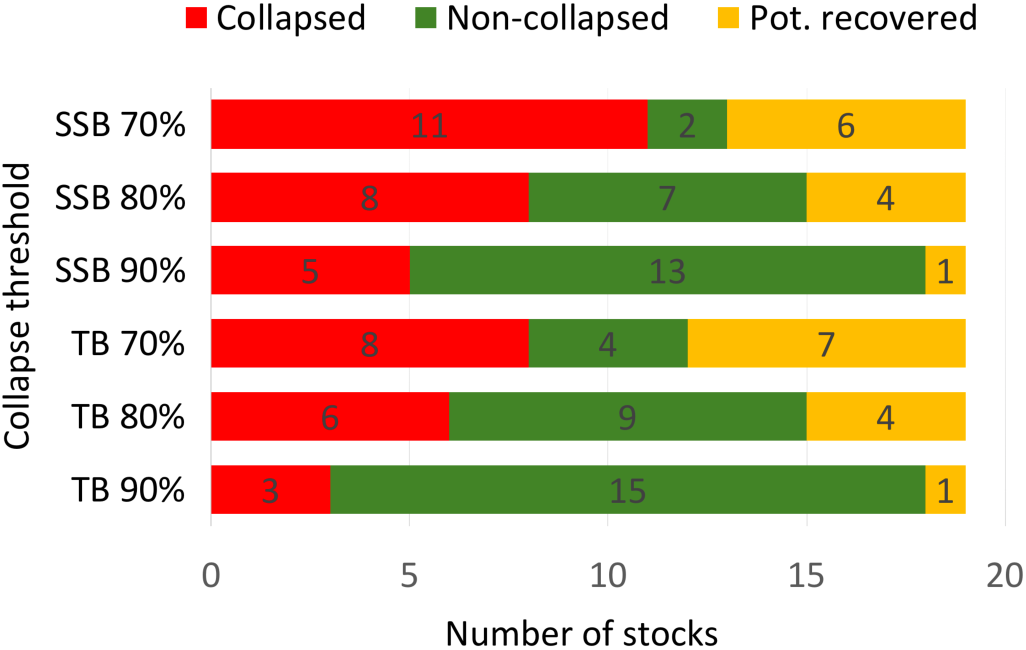
Summary of the analysis results showing the effect of diverse thresholds on the classification of stocks according to the latest data. Numbers inside the bars indicate the number of stocks in each classification. Potentially recovered (“Pot. recovered”) stocks are additional to the collapsed stocks, i.e. if a stock is classified as “potentially recovered”, it has collapsed in the past but has potentially recovered from this state at present.

## DISCUSSION

The proposed definition of collapse will help to avoid the scientific and socioeconomic conflicts that arise from contrasting status assessments, and aid in the allocation of limited management resources to the most severe cases. Distinguishing short-term fluctuations (cases where recruitment is not impaired) from severe collapses can only be made by taking into account the duration of the depletion state. The inclusion of prolonged depletion in the definition of collapse is of great significance because it indicates insufficient management measures, decreased ecosystem resilience, or that the causes that lead to the low recruitment persist (15,24,64,69). The inclusion of recovery potential enables the identification of fish stocks that are recovering from a collapsed state (69), meaning that “collapsed” is no longer an appropriate classification because growth is positive. This is especially important in multispecies fisheries where classifying one species as collapsed (e.g., cod, North Sea) could restrict catches of other fisheries that catch the collapsed species as bycatch (e.g., haddock) (70). Furthermore, few previous definitions were formulated in terms of generation times (or any suitable proxy for population turnover), which ties stock dynamics to biological production and makes a definition robust and comparable to the diverse life histories of marine fishes (65). The downside of this approach is that a generation’s worth of data is required to evaluate any particular year for prolonged depletion. This requires stock-specific estimates of population turnover and also means that the final year(s) cannot be evaluated, but we still believe it is absolutely necessary to distinguish severe collapses from gradual declines and natural fluctuations. In general, data availability will constrain the number of stocks to which our definition is applicable. However, for a substantial set of socioeconomically important stocks, our definition offers the means to standardize collapsed status with consideration for the diverse life-histories of fishes.

A limitation of our definition is that recovery potential is not considered in terms of possible changes in the geographical distribution of the stock (e.g., 71), or its age- or size-structure (c.f., 69). This choice was made because the inclusion of these aspects would involve higher data requirements, severely limiting the number of stocks that can be categorized. Also, the generation of collapse thresholds may be sensitive to interannual fluctuations in abundance due to environmental stochasticity and/or uncertainty in the estimates of abundance (72), for instance, spikes in biomass due to large recruitment events. Although time series smoothing would help to remove fluctuations, a more appropriate approach would be to implement the definition within a Bayesian framework to account for uncertainties in biomass estimates by generating a distributional threshold (see 72). Lastly, fish stock collapse definitions such as those discussed in this paper focus on species-level dynamics only, and exclude the ecosystem context. The inclusion of ecosystem-level signals could reveal additional important aspects of collapse and recovery dynamics (73,74).

We suggest establishing a common fish stock collapse definition to build a foundation for interdisciplinary cooperation as well as a platform for the stringent communication of science and the state of natural resources. Sharing of scientific tools and common data interpretation between the disciplines could reduce differences between conservation scientists and fisheries scientists (67). Finally, avoiding sensationalism (alarmist narratives) is important for a scientific topic of high interest for a wide audience. Carefully chosen narratives may generate greater probability of meaningful action, an example of which is Rachel Carson’s Silent Spring narrative (21). Still, science vocabulary should be based on the contents of the findings rather than the marketability of the findings (20). Our definition can help limit alarmist collapse narratives in science and public media to those cases where fish stock declines actually imply severe risks for fish production and ecosystem functioning. Whilst robust, standardized classifications of stock collapses also prevents scenarios thatunderestimate the severity of the degraded state of a stock, and thus continuing with “business as usual” harvesting strategies.

## CONCLUSIONS

The past two decades have witnessed considerable scientific debate regarding the state of the world’s fish stocks and the occurrence rate of collapses. Here, we have showed that numerous definitions exist to determine whether a fish stock has collapsed, and suggest that the lack of a unified definition has led to contrasting views on the state of fish stocks. We provide an operational definition of fish stock collapse that is based on life history and time series data. We demonstrate that our definition can be used to assess the status of diverse commercially important fish stocks, helping to reconcile contrasting views about the global state of fish stocks, and providing guidance on prioritizing scientific efforts and policy solutions to secure sustainable fisheries. Using the proposed definition in a scientifically consistent way will promote the clear and accurate communication necessary for science to affect fisheries policies. With robust definitions and careful interpretation of findings, co-operation across disciplines can be achieved to secure healthy fish stocks.

## DATA AND CODE AVAILABILITY

Data sources for the time series analysis and collapse definitions are listed in the Supplementary Materials. The complete dataset and computer code (R language) is available upon request.

## SUPPLEMENTARY INFORMATION

Is available in the online version of this paper.

## ACKNOWLEDGEMENTS

We thank the Nordic Centre of Excellence NorMER programme early career scientists for their great contribution towards the literature review and collapse discussions. We would like to thank Jeff Hutchings and Keith Brander for valuable comments on the “collapse” problematics.

## AUTHOR CONTRIBUTIONS

JY and WEB jointly conceived the study idea. TB and NCS supervised the project. JY, WEB, SB, GR, BW, AR and JDW contributed to the literature review. JY, WEB, KHA, SB, ML, MCN, LR, GR, BW, JDW, GO, AR and NCS contributed to stock selection and data collection. WEB performed the analysis. JY, WEB, GO, TB and NCS interpreted the results and discussed the implications with assistance from all authors. JY, WEB, GO, TB and NCS prepared the initial manuscript and all authors contributed to the revisions.

## AUTHOR INFORMATION

This paper is a result of two workshops organized within the Nordic Centre of Excellence NorMER programme. The authors declare no competing interests. Correspondence and requests for materials should be addressed to NCS and TB.

## REFERENCES

1. Garcia SM, Grainger RJR. Gloom and doom? The future of marine capture fisheries. Philos Trans R Soc Lond B Biol Sci. 2005;360(1453):21–46.

2. Pauly D, Hilborn R, Branch TA. Does catch reflect abundance. Nature. 2013;494:3–6.

3. Pauly D, Watson R, Alder J, B PTRS. Global trends in world fisheries: impacts on marine ecosystems and food security. Philos Trans R Soc B Biol Sci. 2005;360(1453):5–12.

4. Jackson JBC, Kirby MX, Berger WH, Bjorndal KA, W BL, Bourque BJ, et al. Historical overfishing and the recent collapse of coastal ecosystems. Science. 2001;293(5530):629–37.

5. Myers RA, Worm B. Rapid worldwide depletion of predatory fish communities. Nature. 2003 May 15;423(6937):280–3.

6. Froese R, Kesner-Reyes K. Impact of fishing on the abundance of marine species. J Mar Sci. 2002;12:1–12.

7. Froese R, Stern-Pirlot A, Kesner-Reyes K. Out of new stocks in 2020: A comment on “Not all fisheries will be collapsed in 2048.” Mar Policy. 2009;33(1):180–1.

8. Worm B, Hilborn R, Baum JK, Branch T a, Collie JS, Costello C, et al. Rebuilding global fisheries. Science (80-). 2009;325(July):578–85.

9. Costello C, Ovando D, Hilborn R, Gaines SD, Deschenes O, Lester SE. Status and solutions for the world’s unassessed fisheries. Science (80-). 2012;338:517–20.

10. Costello C, Ovando D, Clavelle T, Strauss C, Hilborn R, Melnychuk M, et al. Global fishery futures under contrasting management regimes. Proc Natl Acad Sci U S A. 2016;113(18):5125–9.

11. Branch TA. Not all fisheries will be collapsed in 2048. Mar Policy. 2008;32:38–9.

12. Branch TA, Jensen OP, Ricard D, Yimin Y, Hilborn R. Contrasting global trends in marine fishery status obtained from catches and from stock assessments. Conserv Biol. 2011;25(4):777–86.

13. Hilborn R. Reinterpreting the state of fisheries and their management. Ecosystems. 2007 Oct 27;10(8):1362–9.

14. Hoelker F, Baere D, Dörner H, di Natale A, Rätz H-J, Temming A, et al. Comment on “Impacts of Biodiversity Loss on Ocean Ecosystem Services.” Science (80-). 2007;316(5829):1285b–1285b.

15. Worm B, Barbier EB, Beaumont N, Duffy JE, Folke C, Halpern BS, et al. Impacts of biodiversity loss on ocean ecosystem services. Science. 2006 Nov 3;314(5800):787–90.

16. Stokstad E. Détente in the fisheries war. Science (80-). 2009;324(April):170–1.

17. Buchen L. Battling scientists reach consensus on health of global fish stocks. Nature. 2009 Jul 30;

18. Pauly D. Aquacalypse now. New Republic: Environment and Energy. 2009.

19. Hilborn R. Faith-based fisheries. Fisheries. 2006;31(11):554–5.

20. Vinkers CH, Tijdink JK, Otte WM. Use of positive and negative words in scientific PubMed abstracts between 1974 and 2014: retrospective analysis. BMJ. 2015 Dec 14;351:h6467.

21. Nixon B. Slow violence and the environmentalism of the poor. J Commonw Postcolonial Stud. 2006;13.2-14.1(3–4):439–43.

22. Myers RA, Hutchings JA, Barrowman NJ. Why do Fish Stocks Collapse? The Example of Cod in Atlantic Canada. Ecol Appl. 1997;7(1):91–106.

23. Toresen R, Østvedt OJ. Variation in abundance of Norwegian spring-spawning herring (Clupea harengus, Clupeidae) throughout the 20th century and the influence of climatic fluctuations. Fish Fish. 2000 Jul 18;1(3):231–56.

24. Neubauer P, Jensen OP, Hutchings JA, Baum JK. Resilience and recovery of overexploited marine populations. Science (80-). 2013;340(6130):347–9.

25. Golden C. Fall in fish catch threatens human health. Nat Hist. 2016;534:317–20.

26. Close DA, Fitzpatrick MS, Li HW. The Ecological and Cultural Importance of a Species at Risk of Extinction, Pacific Lamprey. Fisheries. 2002 Jul;27(7):19–25.

27. Barnosky AD, Hadly E a., Bascompte J, Berlow EL, Brown JH, Fortelius M, et al. Approaching a state shift in Earth’s biosphere. Nature. 2012;486(7401):52–8.

28. Rockström J, Steffen WL, Noone K, Asa P, Stuart Chapin III FS. Planetary Boundaries: Exploring the safe operating space for humanity. Ecol Soc. 2009;14(2):32.

29. Hughes TP, Barnes ML, Bellwood DR, Cinner JE, Cumming GS, Jackson JBC, et al. Coral reefs in the Anthropocene. Nature. 2017;in press.

30. Reyer CPO, Rammig A, Brouwers N, Langerwisch F. Forest resilience, tipping points and global change processes. Gibson D, editor. J Ecol. 2015 Jan 1;103(1):1–4.

31. Dai L, Vorselen D, Korolev KS, Gore J, Hutchings JA, Reynolds JD, et al. Generic indicators for loss of resilience before a tipping point leading to population collapse. Science. 2012 Jun 1;336(6085):1175–7.

32. Lever JJ, van Nes EH, Scheffer M, Bascompte J. The sudden collapse of pollinator communities. Jordan F, editor. Ecol Lett. 2014 Mar 6;17(3):350–9.

33. Lindenmayer D, Messier C, Sato C. Avoiding ecosystem collapse in managed forest ecosystems. Front Ecol Environ. 2016;14(10).

34. Goossens B, Chikhi L, Ancrenaz M, Lackman-Ancrenaz I, Andau P, Bruford MW. Genetic signature of anthropogenic population collapse in orangutans. Mace G, editor. PLoS Biol. 2006 Feb 24;4(2):e25.

35. Dobson A, Lodge D, Alder J, Cumming GS, Keymer J, McGlade J, et al. Habitat loss, trophic collapse, and the decline of ecosystem services. Ecology. 2006 Aug 11;87(8):1915–24.

36. Connell SD, Fernandes M, Burnell OW, Doubleday ZA, Griffin KJ, Irving AD, et al. Testing for thresholds of ecosystem collapse in seagrass meadows? Conserv Biol. 2017;1–12.

37. Springer AM, Estes JA, van Vliet GB, Williams TM, Doak DF, Danner EM, et al. Sequential megafaunal collapse in the North Pacific Ocean: an ongoing legacy of industrial whaling? Proc Natl Acad Sci U S A. 2003 Oct 14;100(21):12223–8.

38. Scheffer M, Carpenter S, Foley JA, Folke C, Walker B. Catastrophic shifts in ecosystems. Nature. 2001 Oct 11;413(6856):591–6.

39. McCann KS. The diversity-stability debate. Nature. 2000 May 11;405(6783):228–33.

40. MacDougall AS, McCann KS, Gellner G, Turkington R. Diversity loss with persistent human disturbance increases vulnerability to ecosystem collapse. Nature. 2013;494(7435):86–9.

41. Norström A V, Nyström M, Jouffray J, Folke C, Graham NAJ. Guiding coral reef futures in the Anthropocene. 2016;490–8.

42. Rutherford J. Too many boats chasing too few fish: The collapse of the Atlantic groundfish fishery and the avoidance of future collapses through free market environmentalism. Stud by Undergrad Res Guelph. 2008;2(1):11–7.

43. MacKenzie BR, Mosegaard H, Rosenberg AA. Impending collapse of bluefin tuna in the northeast Atlantic and Mediterranean. Conserv Lett. 2009 Feb;2(1):26–35.

44. Hjermann DØ, Ottersen G, Stenseth NC. Competition among fishermen and fish causes the collapse of Barents Sea capelin. Proc Natl Acad Sci U S A. 2004;101:11679–84.

45. Zwolinski JP, Demer D a. A cold oceanographic regime with high exploitation rates in the Northeast Pacific forecasts a collapse of the sardine stock. Proc Natl Acad Sci. 2012;109(11):4175–80.

46. Longo SB, Clark B. The commodification of Bluefin Tuna: the historical transformation of the Mediterranean fishery. J Agrar Chang. 2012;12(2–3):204–26.

47. Schrank WE. The Newfoundland fishery: ten years after the moratorium. Mar Policy. 2005;29(5):407/420.

48. Holt S. Sunken billions - but how many? Fish Res. 2009;97(1–2):3–10.

49. Olsen EM, Heino M, Lilly GR, Morgan MJ, Brattey J, Ernande B, et al. Maturation trends indicative of rapid evolution preceded the collapse of northern cod. Nature. 2004 Apr 29;428(6986):932–5.

50. Litzow M a., Mueter FJ, Daniel Urban J. Rising catch variability preceded historical fisheries collapses in Alaska. Ecol Appl. 2013;23(6):1475–87.

51. ICES. Stock Assessment Database [Internet]. Copenhagen; [cited 2016 May 1]. Available from: http://ices.dk/marine-data/dataset-collections/Pages/default.aspx

52. Ricard D, Minto C, Jensen O, Baum J. Evaluating the knowledge base and status of commercially exploited marine species with the RAM Legacy Stock Assessment Database. Fish Fish. 2013;13(4):380–98.

53. Hill K, Crone PR, Dorval E, Macewicz BJ. Assessment of the Pacific sardine resource in 2015 for U.S. management in 2015-2016. U.S Department of Commerce, La Jolla. 2015.

54. Lindegren M, Checkley DM, Rouyer T, MacCall AD, Stenseth NC. Climate, fishing, and fluctuations of sardine and anchovy in the California Current. Proc Natl Acad Sci U S A. 2013 Aug 13;110(33):13672–7.

55. Bonanomi S, Pellissier L, Therkildsen NO, Hedeholm RB, Retzel A, Meldrup D, et al. Archived DNA reveals fisheries and climate induced collapse of a major fishery. Sci Rep. 2015 Dec 22;5(1):15395.

56. Hutchings JA. Collapse and recovery of marine fishes. Nature. 2000;406(August):882–5.

57. Rainer F, Gianpaolo C, Kristin K, Nazli D. Revisiting safe biological limits in fisheries. Fish Fish. 2016 Feb 16;17(1):193–209.

58. Froese R, Coro G, Kleisner K, Demirel N. Estimating fisheries reference points from catch and resilience. Fish Fish. 2017 Apr 27;18(3):506–26.

59. Pauly D. Anecdotes and the shifting baseline syndrome of fisheries. Trends Ecol Evol. 1995;10(10):430.

60. Oxford University Press. English Oxford Living Dictionaries [Internet]. 2017 [cited 2017 Jul 13]. Available from: https://en.oxforddictionaries.com

61. Taylor MS. Innis Lecture?: Environmental crises?: past, present, and future. 2009;42(4):1240–75.

62. Mullon C, Fréon P, Cury P. The dynamics of collapse in world fisheries. Fish Fish. 2005;6:111–20.

63. IUCN. IUCN Red List Categories and Criteria. Version 3.1. Second edition. Gland, Switzerland and Cambridge, UK: IUCN; 2012. iv + 32.

64. Hutchings JA, Reynolds JD. Marine fish population collapses: consequences for recovery and extinction risk. Bioscience. 2004;54(4):297.

65. Reynolds JD, Dulvy NK, Goodwin NB, Hutchings JA. Biology of extinction risk in marine fishes. Proc R Soc B Biol Sci. 2005;272(1579):2337–44.

66. Hiers JK, Jackson ST, Hobbs RJ, Bernhardt ES, Valentine LE. The precision problem in conservation and restoration. Trends Ecol Evol. 2016;31(11):820–30.

67. Salomon AK, Gaichas SK, Jensen OP, Agostini VN, Sloan NA, Rice J, et al. Bridging the divide between fisheries and marine conservation science. Bull Mar Sci. 2011;87(2):251–74.

68. Fernandes PG, Cook RM. Reversal of fish stock decline in the northeast atlantic. Curr Biol. 2013;23(15):1432–7.

69. Lotze HK, Coll M, Magera AM, Ward-Paige C, Airoldi L. Recovery of marine animal populations and ecosystems. Trends Ecol Evol. 2011;26(11):595–605.

70. Burgess MG, Polasky S, Tilman D. Predicting overfishing and extinction threats in multispecies fisheries. Proc Natl Acad Sci. 2013 Oct 1;110(40):15943–8.

71. Pinsky ML, Worm B, Fogarty MJ, Sarmiento JL, Levin SA. Marine Taxa Track Local Climate Velocities. Science (80-). 2013;341(6151):1239–42.

72. Aagaard K, Lockwood JL, Green EJ. A Bayesian approach for characterizing uncertainty in declaring a population collapse. Ecol Modell. 2016;328:78–84.

73. Pikitch EK, Santora C, Babcock E a, Bakun A, Bonfil R, Conover DO, et al. Ecosystem-based fishery management. Science (80-). 2004;305(February):346–7.

74. Pedersen EJ, Thompson PL, Ball RA, Fortin M, Tarik C, Link H, et al. Signatures of the collapse and incipient recovery of an overexploited marine ecosystem. 2017;1–31.

